# Population-level structural variant characterization from pangenome graph

**DOI:** 10.1101/2025.07.06.663386

**Authors:** Songbo Wang, Tun Xu, Pengyu Zhang, Kai Ye

**Affiliations:** School of Automation Science and Engineering, Faculty of Electronic and Information Engineering, Xi’an Jiaotong University, Xi’an, China; MOE Key Lab for Intelligent Networks & Networks Security, Faculty of Electronic and Information Engineering, Xi’an Jiaotong University, Xi’an, China; Department of Gynecology and Obstetrics, Center for Mathematical Medical, The First Affiliated Hospital of Xi’an Jiaotong University, Xi’an, China; Genome Institute, The First Affiliated Hospital of Xi’an Jiaotong University, Xi’an, China; Faculty of Science, Leiden University, Leiden, The Netherlands

## Abstract

Population-level structural variant (SV) profiling is crucial in the era of pangenomes. However, identifying SVs from genome assemblies and pangenome graphs remains a significant challenge. Here we present Swave, a sequence-to-image, deep-learning based method that accurately resolves both simple and complex SVs, along with their population characteristics, from assembly-derived pangenome graphs. Swave introduces ‘projection waves’ to summarize the dotplot images that capture mapping patterns between reference and SV-indicating alleles in pangenome. These images are analyzed by a recurrent neural network to distinguishes true SV signals from background noise introduced by genomic repeats. Swave demonstrates superior performance in both SV type classification and genotyping compared to existing methods. When applied to a healthy cohort (n=334) and a rare-disease cohort (n=574), Swave reveals complex and polymorphic SV patterns across human populations and identifies potentially pathogenic SVs. These advancements will facilitate the creation of comprehensive population-level SV catalogs, deepening our understanding of SVs in genetic diversity and disease associations.

## Introduction

Structural variants (SV) are genomic alterations larger than 50 base-pairs (bp), categorized into simple SVs^1^ (SSVs, e.g., insertions, deletions inversions and duplications) and complex SVs (CSVs) with more than one internal breakpoints or subcomponents^1,2^. Recent advances in whole genome sequencing have underscored the critical roles of SV in development^3^, genetic disorders^4^, and cancers^5^. The emergence of long-read sequencing (LRS) technologies has markedly improved SV discovery^1^, boosting the number of detectable SVs by 2-4.25 folds^6^, and enhanced resolution of structural complexity, particularly for CSVs. Several LRS-based SV callers have been developed^2,7–11^ using either model-matching or deep-learning strategies^12^. Concurrently, improvements in genome assembly methods have resulted in high-quality genome assemblies that offer longer, more precise sequences, outperforming read-based SV detection^13,14^. However, existing assembly-based SV callers remain limited by model-matching approaches^15,16^, which restrict detection to known SV types and often overlook uncharacterized or complex SVs.

With the declining cost of genome sequencing and computing, population-level SV analysis using assemblies is now feasible^17–19^. Examining SVs across large cohorts enables comprehensive characterization of SV landscapes and population-specific features, offering insights into human evolution and clinical interpretation^20,21^. Several gene loci (e.g. *AMY1*^22,23^, *MUC5B*^24^) have been examined at population-level, revealing contributions of SVs to natural selection and adaptation, highlighting the need for automated, genome-wide screening of evolution-associated SV loci. Clinically, distinguishing pathogenic SVs from benign ones observed in health populations, requires a robust SV reference derived from population-scale datasets.

A critical component of population-level SV analysis is cross-sample merging, which integrates individual SV callsets into population profiles reflecting SV genotypes and frequencies^25–27^. However, current merging techniques, primarily based on SV region overlap and sequence similarity, are prone to false positives and missing genotypes^2,28^. They struggle in genome repetitive regions, where mapping ambiguities causing variable SV lengths, and in CSV regions, where complex structures with nested breakpoints are often misclassified or missed entirely.

Pangenome graph, which compactly represents multiple genomes along with their similarities and differences, is well suited to population-scale SV applications^17–19,29,30^. Through careful alignments^31–34^, graph snarls encode SV alleles within nodes and paths^18,35,36^. For any given assembly, pangenome graph construction tools annotate its specific path and SV allele within each snarl, facilitating SV genotyping. Still, classifying SV alleles in pangenome graph remains a bottleneck. Existing LRS- or assembly-based SV calling methods, whether model-based or deep-learning-based, are not designed for pangenome graphs. Instead, current tools rely on length differences between reference and alternative alleles to identify deletions and insertions^29,30,37^, which is insufficient for capturing the full landscape of SV diversity in populations.

To address these limitations, we propose Swave, a method that leverages assemblies and pangenome graphs to enable SV discovery from individual genomes to population-wide datasets. Swave parses SV-indicating alleles from pangenome graphs and realign them against reference alleles to generate base-level dotplot images. These images are then transformed into projection waves that summarize alignment conditions at each genomic location. By comparing against the waves generated from reference genome backgrounds, Swave reduces the disruptions from genome repetitive sequence. A recurrent neural network (RNN) is then applied to classify SV types. Benchmarking against state-of-the-art methods, Swave exhibits superior accuracy in both SV classification and genotyping using assembly-derived pangenomes. Applied to healthy cohorts (334 haplotypes), Swave reveals the population-level complexity and polymorphism of challenging SSVs (inversions) and previously underestimated CSVs, especially rare ones with population allele frequency (AF) below 1%. In a rare disease cohort comprising 287 proband genomes (574 haplotypes), Swave identifies potentially pathogenic SVs characterized by singleton and exon-disrupting events, including 888 SSVs and 34 CSVs. These findings broaden the spectrum of pathogenic variants from small-scale mutations (e.g., SNPs) to SVs that can induce more extensive genomic damage.

## Results

### Algorithm overview

Swave comprises three key modules:

1. **Graph module for SV allele deconstruction (Fig.1a, Extended Data Fig. 1** and **Methods)**. Using both reference genome and sample assemblies, Swave constructs pangenome graph using Minigraph’s incremental graph generation^31^. Within these graphs, graph snarls (graph substructures indicating local variation) are considered as candidate SV loci. Paths traversing distinct sequence nodes within each snarl are extracted as SV allele sequences. For each SV allele, Swave determines the carrier sample(s) and calculates its frequency against the whole snarl using the outputs of Minigraph ‘call’ command. The precise classification of these alleles into SV types is deferred to the next two modules.
2. **Sequence-to-image module for allele realignment and resolution (Fig.1b, Extended Data Fig. 2** and **Methods)**. To classify each SV allele sequence (ALT), Swave performs elaborate realignment against the reference sequence (REF) using a dotplot (REF2ALT dotplot). In this image-based representation, matched and reversely-matched kmers are denoted as black dots. While dotplots sensitively reflect SV-induced sequence differences, they are also cluttered by noise signals from genome repetitive sequences, severely disturbing the identification of SVs. To address this, Swave introduces two dotplot waves by projecting the dotplots to the REF axis that quantify dot density for both matched and reversely-matched orientations. Then we realign REF sequence against itself to obtain a REF2REF dotplot and use its projected waves as genomic background to reduce the noises from repetitive sequence. Briefly, non-SV regions in the REF2ALT dotplot mirror the background REF2REF waves, while true SVs produce characteristic wave transformations relative to the REF2REF background waves depending on the SV type. Therefore, the SV types are implied from the comparison between the REF2REF and REF2ALT waves.
3. **Deep learning module for SV types prediction (Fig.1c, Extended Data Fig. 3** and **Methods).** Swave employs an RNN to automatically classify the wave differences into SV types. A simulated dataset is used to train a Bidirectional Long Short-Term Memory (Bi-LSTM) network. Every recurrent step receives the wave difference at a reference position and output a predicted SV type among five canonical SV types: insertion, deletion, inversion, duplication, and duplicated inversion. Bidirectional recurrences enable the model to incorporate both upstream and downstream sequence contexts for accurate classification. Briefly, duplications exhibit increased signal intensity, deletions show diminished signals, inversions present reversed patterns, and duplicated inversions combine both features of duplication and inversion. Consecutive positions predicted to share the same SV type are merged into single SV components. When multiple such components shared breakpoints within a dotplot, Swave will sequentially merge them into a CSV type

### Accurate SV detection

#### Performance Evaluation

We assessed the performance of Swave alongside other assembly-based and pangenome-based SV calling approaches using both simulated and published assemblies. Assembly-based callers, including PAV^15^ and SVIM-asm^16^, were paired with three SV merging tools (SURVIVOR^25^, Jasmine^26^ and Truvari^27^) to integrate single-sample callsets into a population-level set. There were further refined using PanPop PART^28^. Pangenome-based SV calling approaches generally classify SVs based on allele length difference within snarls. Simple and bi-allelic snarls were directly classified into deletions or insertions based on allele length differences. In contrast, complex and multi-allelic snarls were firstly processed to reduce allele complexity before classification. In this evaluation, we applied a widely adopted pipeline in recent pangenome studies (e.g., HPRC^17^ and CPC^18^) where the pangenome graphs were processed sequentially using vg^35^, vcfbub and vcfwave^37^ (vg-vcfwave for short) for SV discovery.

##### Individual-level evaluation

Using the HG002 assembly and Tire 1 high-confidence SVs^38^ as ground truth (**Fig. 2a and Supplementary Table 1**), Swave achieved the highest F1-score (0.957), outperforming the two assembly-based callers, PAV (0.947) and SVIM-asm (0.951). Vg-vcfwave lagged behind with an F1-score of 0.791, likely due to its reliance on allele length differences alone for SV classification. For individual-level CSV calling, Swave was compared to SVision-pro^2^ on its previously simulated CSV ground truth (**Fig. 2b and Supplementary Table 1**). Swave achieved an F1-score of 0.956, substantially higher than SVision-pro on assemblies (0.655) and nearly matched its performance on long-reads (0.967). This reflects Swave’s specialized design for assemblies, while SVision-pro was optimized for read-based input.

**Fig. 1.**
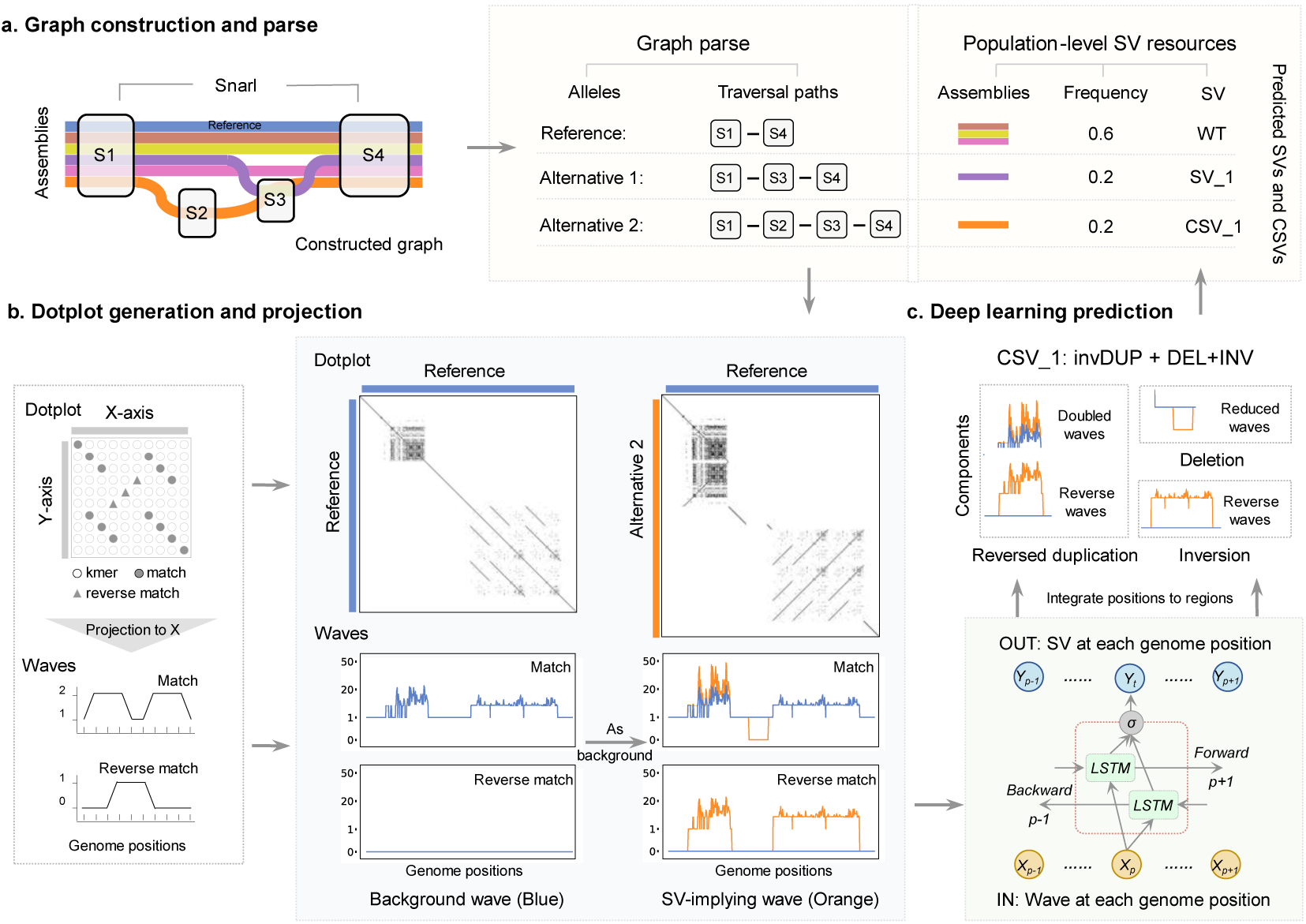
Schematic overview of Swave. **a, Pangenome graph construction and allele extraction.** Swave extracts allele paths for both reference and individual assemblies from a pangenome graph. **b, Sequence-to-image transformation**. Dotplot images depicting structural differences between reference sequence and alternative allele sequences are projected into waves to extract both background and SV-indicating signals. **c, Deep learning-based SV classification**. A RNN takes in the wave signals and assigns SV types based on learned sequence-context patterns.

**Fig. 2.**
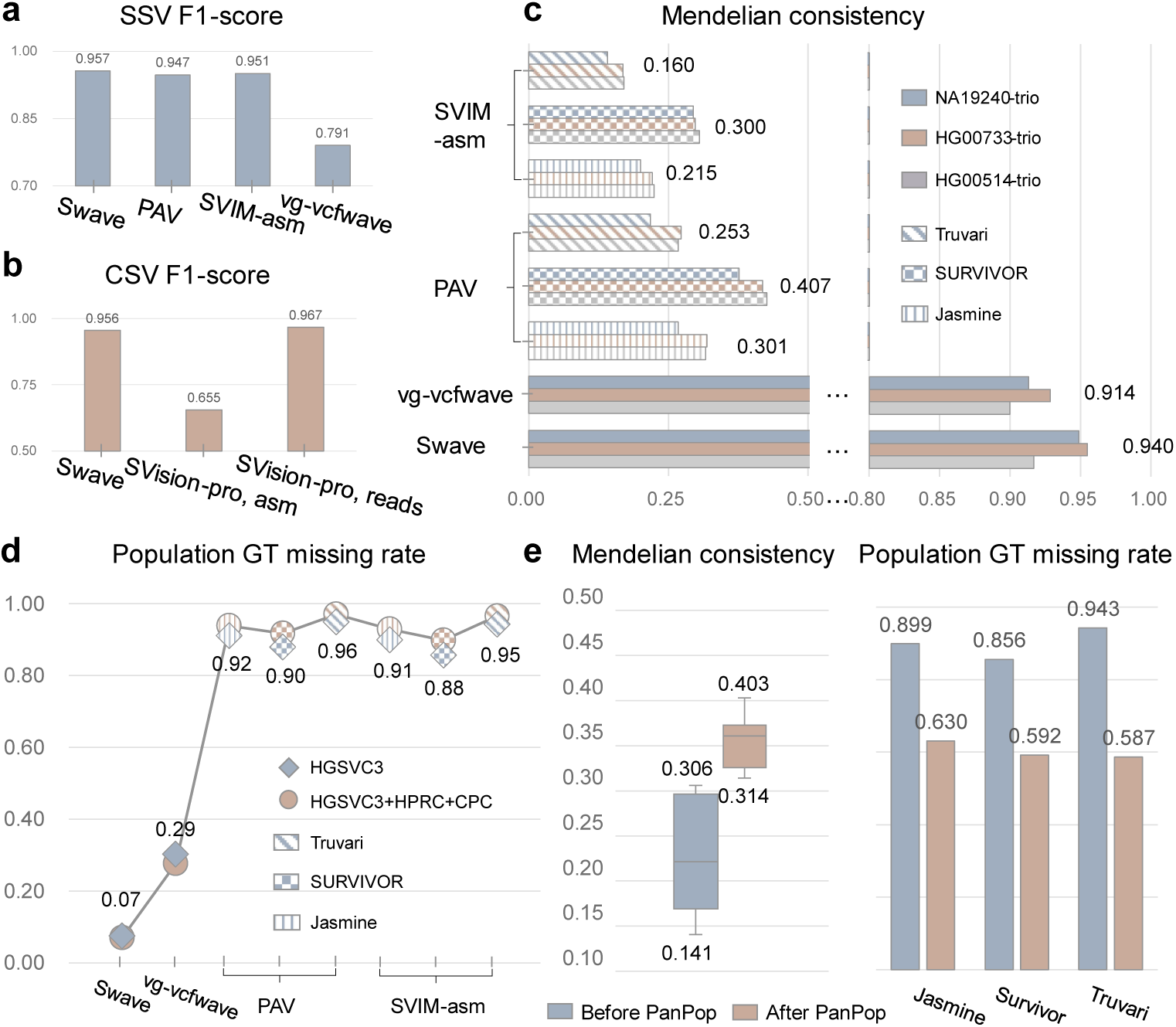
Benchmarking the Performance of Swave and comparative methods. **a,** F1-socre comparison for simple structural variant (SSV) detection. **b,** F1-score comparison for complex structural variant (CSV) detection. **c,** Mendelian consistency comparison in tree parent-child trios. Mean consistency values are indicated to the right of each bar. **d,** Genotyping (GT) missing rate across two population datasets. Average missing rates on the two datasets are noted on the plot. **e,** Improvements in genotyping performance following PanPop refinement. Boxplots represent the interquartile range (IQR) from first (Q1) to third (Q3) quartiles. Whiskers extend to values within Q1 − 1.5*IQR and Q3 + 1.5*IQR. Minimum and maximum values are indicated.

##### Trio- and population-level evaluation

Pangenome-based methods, like Swave and vg-vcfwave, can directly output genotype-aware population callset, whereas assembly-based callers depend on downstream merging procedure to aggregate individual callsets. We evaluated performance on three parent-child trio assemblies (CHS, PUR and YRI; **Fig. 2c and Supplementary Table 2**). Swave achieved the highest mendelian consistency (mean 0.940), surpassing vg-vcfwave (0.914) and merging-based assembly approaches (0.160-0.407). At the population-scale, we assessed genotyping completeness using the HGSVC assemblies^19^ (comprising 130 haplotypes, **Fig. 2d and Supplementary Table 3**). Merged callsets from assemblies exhibited high genotype missing rates (0.856-0.949). Vg-vcfwave reduced this to 0.303 while Swave further minimized the missing rate to 0.075. Notably, when scaled to larger cohorts (adding HPRC and CPC assemblies), Swave and vg-vcfwave maintained stable performance while merging-based approaches exhibited worsening missing rate (0.898-0.970), restricting their utility for downstream population SV analysis. PanPop PART improved the performance of merging-based approaches slightly (Mendelian consistency: 0.332-0.381; genotyping missing rate: 0.587-0.630, **Fig. 2e and Supplementary Table 2-3**), but still underperformed than Swave. Overall, current pangenome and assembly-based tools struggle with either SV classification or multi-sample genotyping—challenges that Swave addresses jointly across SSVs and CSVs.

#### Enhanced resolution of large and complex inversions

Large inversion polymorphisms and their complex patterns have been associated with genomic instability and genetic disorders^39,40^, yet automated detection of them is still challenging for current methods. Since neither the GIAB nor the simulated ground truth callset included large inversions, we benchmarked Swave’s inversion-calling performance using published calls from the HGSVC dataset (130 haplotypes, **Fig.3a**, **Methods**). Swave identified 156 inversion snarls encompassing 322 alleles, including 129 balanced inversions and 193 complex inversions (**Supplementary Table 4**). For comparison, the latest PAV calls on the same assemblies reported 189 balanced inversions, while SVIM-asm detected only 18 balanced inversions, and was therefore excluded from downstream analysis. Given that Swave reported notably more inversions compared to other approaches, we performed computational validations. First, we mapped the reconstructed inversion-feature sequences back to their carrier assemblies, achieving an average mapping integrity of 99% (**Extended Data Fig.4a and Supplementary Table 5**), indicating that these inversion-feature sequences could be forwardly and continuously aligned without clipping. Next, we applied two validation metrics sourced from TT-Mars^41^ and Vapor^42^, confirming that 99% of Swave’s inversion calls were supported by at least one validation metric and 97% were supported by both (**Extended Data Fig.4b and Supplementary Table 5**).

##### Balanced inversion

Balanced inversion refers to a directly inverted sequence without any loss or gain of sequence (**Fig.3b**). BEDtools^43^ intersection revealed that 117 of 189 balanced inversions from PAV calls overlapped with Swave balanced calls, while 61 with Swave complex inversions (**Supplementary Table 6**). Strikingly, among these 117 overlapped inversion calls, Swave consistently reported shorter inversion lengths than PAV (**Fig.3b**). We validated this against a high-confidence inversion benchmark compiled from assemblies, Strand-seq, Bionano and manual curation^39^. Swave’s length estimates showed tight concordance with this benchmark (**Fig.3c and Supplementary Table 7**, *Wilcoxon rank sum test*, *p-value* = 2.3e-19), yielding a Pearson correlation of 0.99 (*p-value* = 6e-72, **Extended Data Fig.4c**), while PAV’s calls were poorly correlated with Pearson correlation of 0.10 (*p-value* = 0.44). During this comparison, we found that 97 out of 117 overlapped balanced inversions were flanked by inverted segmental duplications (SDs, **Supplementary Table 8**), consistent with previous studies linking large inversions with recurrent SDs. These SDs likely misled PAV’s breakpoint resolution, leading to systematic overestimation of inversion lengths (**Extended Data Fig.4d**).

##### Complex and polymorphic inversions

Of the 156 inversion-related snarls identified by Swave, approximately two-thirds (n=128, **Fig. 3d** and **Supplementary Table 4**) corresponded to flanked events, where inversions (n=43) or duplicated inversions (n=85) were neighbored by additional SV breakpoints precisely at their boundaries. Notably, Swave uncovered a previously undescribed subclass of complex inversion, termed as scarred inversions (n=71, **Fig.3d-e and Supplementary Table 9**), defined by insertion or deletion breakpoints (‘scars’) occurring inside the inversion bodies. Most scarred inversions (63/71, **Supplementary Table 9**) contained a single internal scar, while seven had two and one included up to four (**Extended Data Fig. 5a**), totaling 81 internal scars. These scars were typically small (**Extended Data Fig. 5b**): 75 of 81 were under 5,000 bp, with the largest spanning 18,451 bp and removing 24% of the original inverted sequence. Existing tools failed to capture the full structure of these scarred inversions (**Fig.3f**). Specifically, PAV misclassified 72% as simple balanced inversions, reported the internal scars in 5%, and entirely missed the remaining 23%. SVIM-asm exhibited the opposite tendency, reporting the scars for 70% of scarred inversions while missing the inversion itself.

**Fig. 3.**
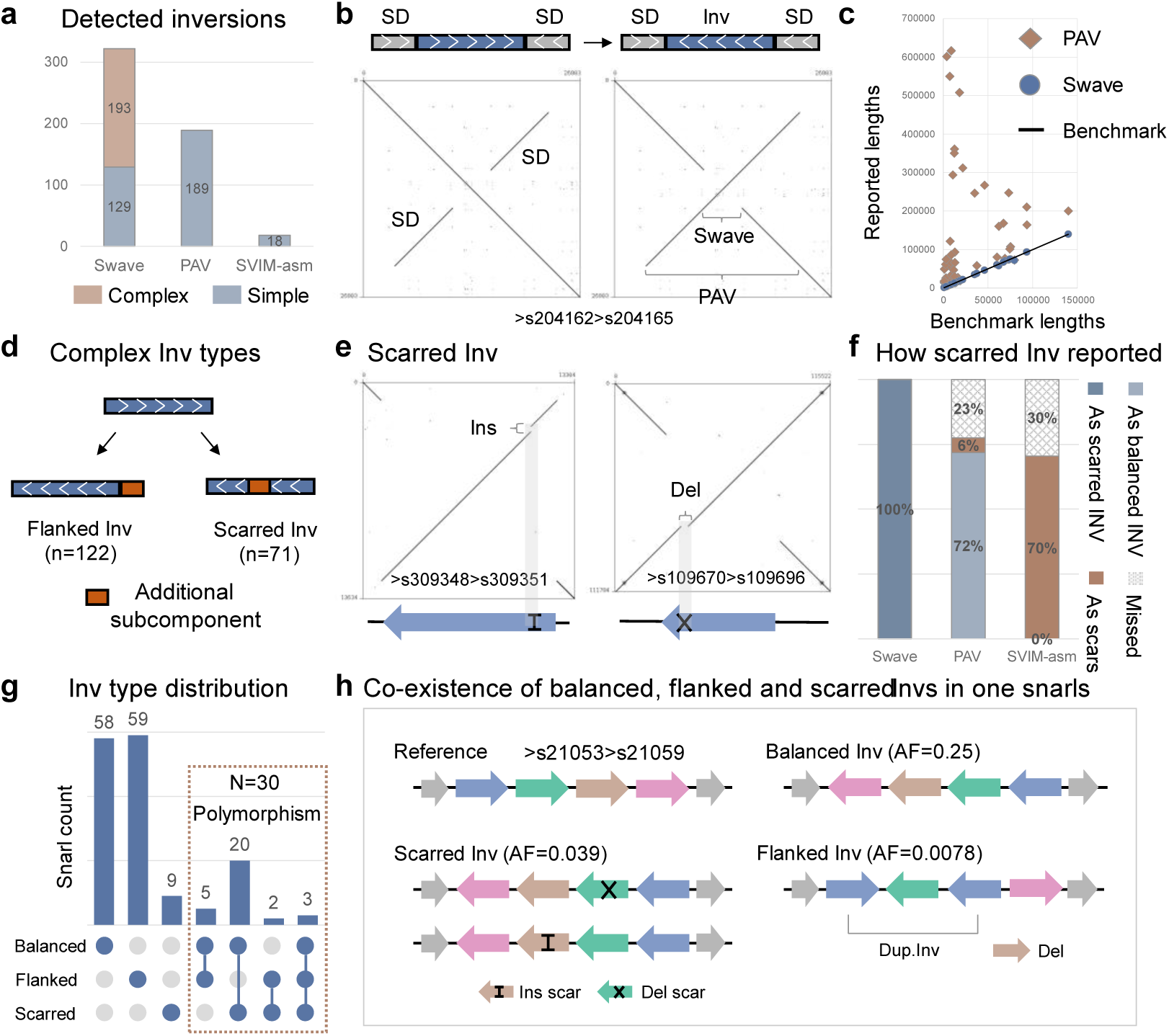
Inversion discovery. **a,** Total numbers of detected inversions from Swave, PAV and SVIM-asm. Only Swave identified complex inversion types. **b,** Example of a balanced inversion (snarl >s204162>s204165) located between inverted segmental duplications (SDs), potentially leading to overestimation of event length. **c,** Comparison of length discrepancies in balanced inversions called by PAV and Swave relative to the benchmark lengths. **d,** Classification of complex inversions detected by Swave into flanked and scarred subtypes. **e,** Representative scarred inversions containing internal insertion scar (left, >s309348>s309351) and deletion scar (right, >s109670>s109696), respectively. **f,** Detection performance for scarred inversions by PAV and SVIM-asm. Neither method successfully resolved these variants. **g,** Distribution of inversion types across the 156 inversion-associated snarls, with 30 snarls containing multiple inversion types. **h,** Example of a multi-category snarl (>s21053>s21059), harboring balanced (AF=0.25), scarred (AF=0.039), and flanked (0.0078) inversion alleles.

Among the 156 inversion-related snarls, majority of them (102, 65%) were bi-allelic (one wide-type allele and one inversion allele) while 54 (35%, **Supplementary Table 4**) were multi-allelic, harboring an average of 4.0 distinct alleles. This high allelic diversity reflects the extensive polymorphism of complex inversions across samples. One form of polymorphism in complex inversions was driven by the diversity in types and lengths of additional breakpoints. In scarred inversions, much of the diversity was attributable to variation in the types and lengths of internal scars (**Extended Data Fig. 5**). For instance, a single graph snarl (>s142244>s142248) harbored five co-existing scarred inversions formed by different combinations of four scar regions (**Extended Data Fig. 5a**). Repeat elements further contributed to this diversity: 40 of 81 scars occurred within repetitive regions (**Extended Data Fig. 5c** and **Supplementary Table 9**), where expansions or contractions likely gave rise to insertion or deletion scars, respectively (**Extended Data Fig. 5d)**. Another form of polymorphism involved the co-occurrence of the three inversion categories (balanced, flanked, and scarred inversions, **Fig.3h**) at individual loci. Of the 54 multi-allelic snarls, 30 featured two inversion classes, and three contained all three (**Fig.3h**). For example, the snarl >s21053>s21059, spanning chr1:247,955,488–247,975,948, included balanced (AF = 0.25), scarred (AF = 0.039) and flanked (AF = 0.0078) forms (**Fig.3i**). The balanced inversion represented the predominant allele, while scarred and flanked alleles introduced unique breakpoint complexities at the same genome locus. A similar pattern was observed in 28 out of the 30 the multi-category inversion snarls featuring a balanced inversion allele, where the balanced form was most common in 18 cases (**Fig.3h**).

### Resolved SVs in Healthy Population

Having demonstrated Swave’s superior performance in detecting of SSVs and CSVs using pangenomes, we next applied it to the pangenomes from three largest healthy cohorts to date: HGSVC (130 haplotypes), HPRC (88 haplotypes) and CPC (116 haplotypes), exploring the population-level profiles of challenging SSVs and previously underestimated CSVs **(Fig. 4a)**. The resulted callset could be found in the **Data Availability**.

**Fig. 4.**
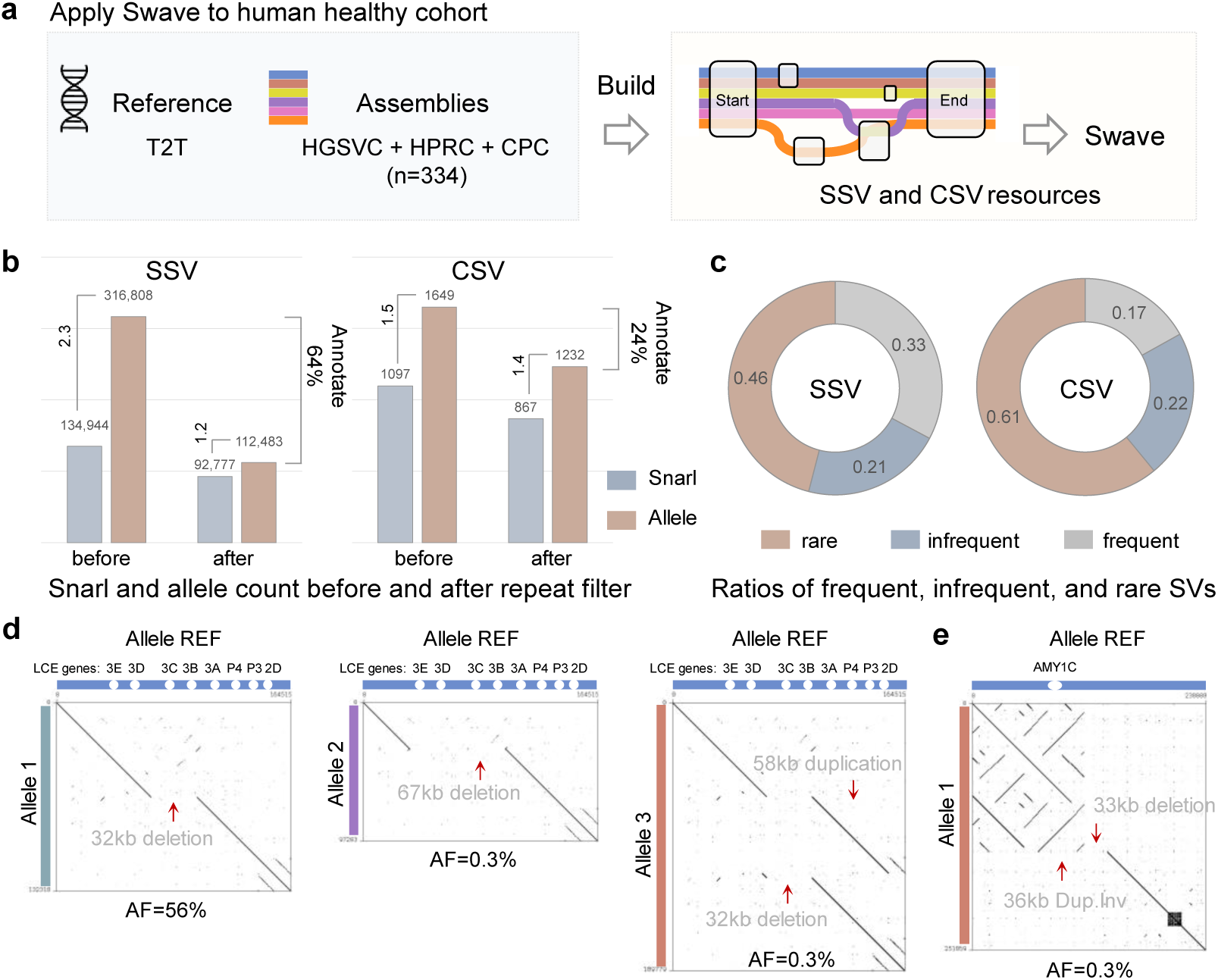
Population-scale discovery of simple and complex structural variants using Swave. **a,** Schematic of Swave’s application to healthy genome cohorts (HGSVC, HPRC and CPC). **b,** Total counts of simple structural variants (SSVs) and complex structural variants (CSVs in the combined HGSVC, HPRC, and CPC cohorts. Repeat annotation was applied to both variant types. Allele counts per snarl are shown to the left of each bar, and the fraction of repeat-overlapping snarls is noted to the right. **c,** The allele frequency distribution of retained SSVs and CSVs, stratified into frequent (AF > 5%), infrequent, or rare (AF < 1%) categories. **d,** A frequent SSV (left: deletion of *LCE3B*/*LCE3C*) and two rare alleles (middle: SSV; right: CSV involving a duplication flanked by a deletion) at the same locus. The rare alleles introduce greater disruption to the *LCE* gene cluster. **e,** Example of a novel and rare CSV locus (a duplicated inversion flanked by a deletion) that increases copy number of *AMY1C*, a gene with known dosage variability.

As a result, 134,944 SSV snarls comprising in total of 316,808 alleles were identified **(Fig. 4b, left)**. Repetitive genomic regions were a major source of multi-allelic SSV snarls. Annotating snarls with over 80% of their variant sequences were repetitive reduced the callset to 92,777 snarls and 112,483 alleles, bringing the average allele number per snarl down from 2.3 to 1.2. Swave also detected 1,649 CSV alleles from 1,097 snarls, of which only 230 were annotated as repetitive. Repetitive sequences had a notably minor impact on CSVs than on SSVs: annotation decreased the average allele count modestly from 1.5 to 1.4, and only 25% of CSVs were annotated, substantially lower than the 64% observed for SSV (**Fig. 4b, right**). The remaining 867 non-repetitive snarls comprised 1,232 complex alleles (**Fig. 4c**). Among these, rare CSVs (AF < 1%, n=755, 61%) were more prevalent than infrequent (1-5%, n=274, 22%) and frequent (>5%, n=203, 17%) variants. Notably, the proportion of rare CSVs exceed that of SSVs (46% vs 61%, **Fig. 4c** and **Supplementary Table 10**), highlighting the need to better characterize these overlooked rare events.

#### Rare CSV alleles

Swave captured rare CSV alleles at both multi-allelic (**Fig. 4d**) and bi-allelic genomic regions (**Fig. 4e**). 237 (31%) of these rare alleles were identified within mixed snarls (n=103) containing also SSV alleles (**Supplementary Table 11**). Among these, 70 snarls featured more common SSVs than CSVs, indicating that the CSVs were minor alleles that either introducing new breakpoints atop existing SSVs (**Fig. 4d**) or restructuring this locus entirely (**Extended Data Fig. 6d**). For example, the well-characterized 32kb deletion affecting *LCE3B*/*LCE3C* genes (**Fig. 4d left**), which was strongly associated with psoriasis and estimated to have persisted for at least 45,000 years^44^, was found alongside two novel singleton alleles (**Extended Data Fig. 6a**). One extended the deletion to 67kb (**Fig. 4d middle**), and the other introduced a 58kb downstream duplication precisely flanking the ancestral 32kb deletion (**Fig. 4d right** and **Extended Data Fig. 6b**). Another notable case occurred upstream of *VIPR2*, a gene linked to schizophrenia risk^45,46^. This region exhibited highly variable repeat expansions/contractions across populations without affecting the gene sequence itself, yet Swave identified a rare 28 kb scarred inversion that partially disrupted the *VIPR2* gene body (**Extended Data Fig. 6d**).

The remaining 518 rare alleles (69%) were from bi-allelic CSV snarls (n=436 and **Supplementary Table 11**) containing only a wild-type allele alongside the CSV allele. Although these snarls occurred outside of other loci, they could still influence the same gene by introducing novel CSV loci. For example, a novel and rare CSV locus (AF=0.3%, **Fig. 4e**) on *AMY1C* gene, known for highly variable copy number driven by palindromic and tandem repeats^22,23^. This novel event occurred downstream of the repeats rather than within it as commonly observed (**Extended Data Fig. 6c**). The variant consisted of a 36kb duplicated-inversion, adding one *AMY1C* copy, that replaced a 33kb sequence located downstream of the repeat boundary.

### Resolved SVs in Rare-Disease Cohort

Identifying pathogenic SVs in rare genetic diseases requires careful exclusion of those found in heathy cohorts, which are typically considered as benign, to pinpoint SVs specifically presented in disease genomes. A recent effort constructed the largest publicly available rare disease pangenome to date, comprising 287 pediatric disease genomes (574 haplotypes) from the Genomic Answers for Kids (GA4K)^21^, alongside 94 HPRC haplotypes as healthy control genomes. Here, we applied Swave to this pangenome to profile SVs and uncover novel complex patterns potentially associated with disease **(Fig. 5a).**

**Fig. 5.**
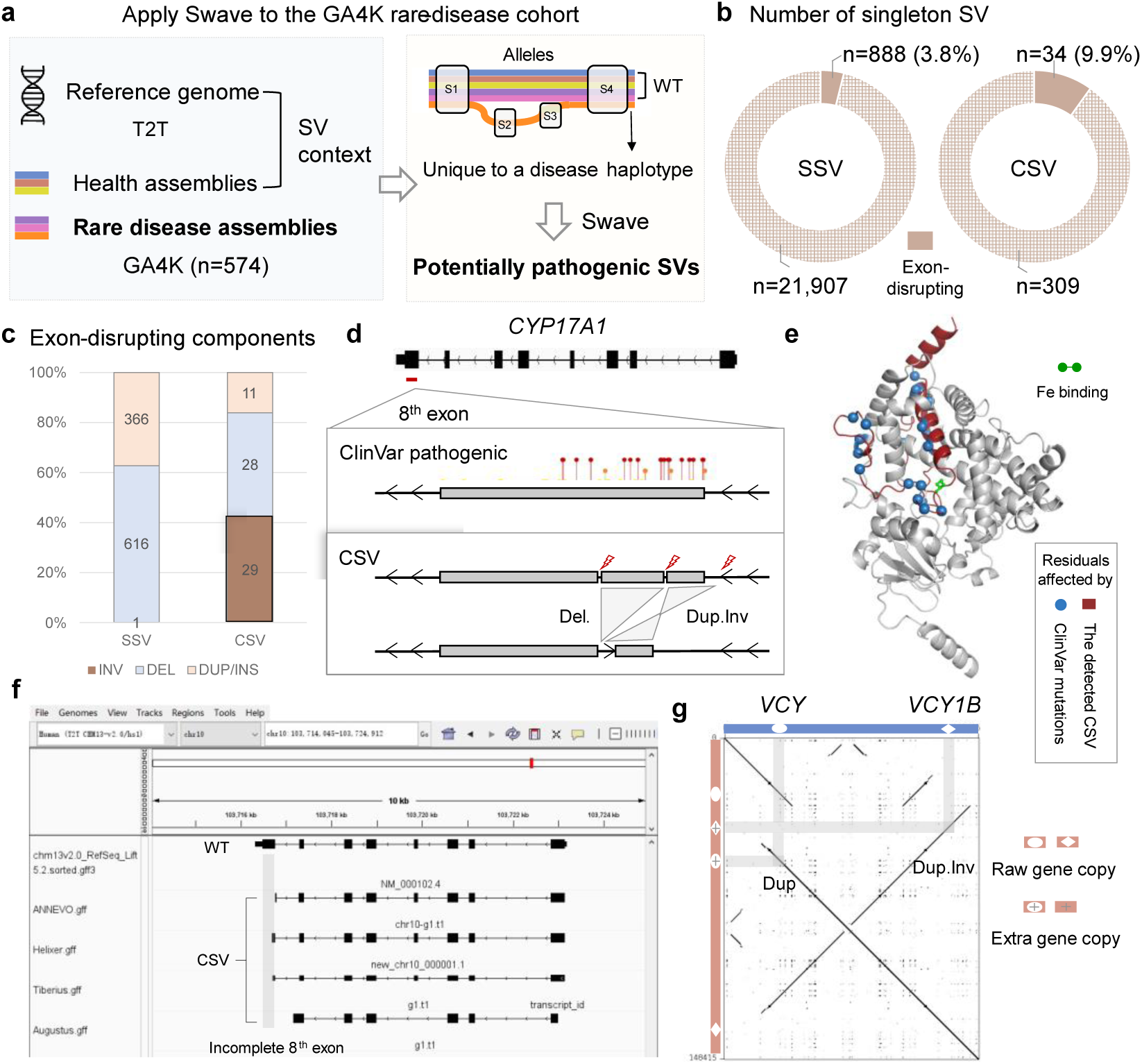
Discovery of potentially pathogenic structural variants in the GA4K rare disease cohort. **a,** Schematic of Swave’s application to the GA4K rare disease cohort. Assemblies from both healthy and rare-disease cohorts were combined to construct a pangenome graph. Alleles exclusively to a single disease genome (singletons) were considered potentially pathogenic variants. **b,** Counts of singleton SSVs and CSVs. Exon-disrupting events are indicated as solid pie segments. The proportion of exon-disrupting CSVs was more than twice that of SSVs. **c,** Component type distribution among exon-disrupting singleton SSVs and CSVs. Exon-disrupting inversions were more frequently observed as CSV subcomponents (n=29) than as standalone SSVs (n=1). **d,** A singleton CSV (a duplicated inversion flanked by a deletion) disrupting the 8^th^ exon of *CYP17A1* which harbors multiple known pathogenic (red pushpins) and likely pathogenic (short orange pushpins) small variants in ClinVar. **e,** Mapping of residue-level disruptions caused by ClinVar pathogenic variants and the CSV detected by Swave. **f,** Comparison of predicted gene models from four genome annotation tools against the raw annotation in the T2T reference genome reveals incomplete structure of 8^th^ exon. **g,** Example of a CSV (a duplicated-inversion flanked by as duplication) that doubled the copy number of two paralogous genes *VCY* and *VCY1B* through distinct subcomponents.

Swave identified 343 singleton CSVs unique to GA4K genomes **(Fig. 5b** and **Supplementary Table 12)**. These variants, supported by only a single disease haplotype and absent from all healthy and other disease haplotypes, were predominantly complex inversions (n = 307; 90%). Although there were far fewer singleton CSVs than singleton SSVs (n=22,795, **Supplementary Table 13**), the proportion of exon-disrupting CSVs (n=34, 9.9%, **Supplementary Table 12**) was more than twice that of SSVs (n=888, 3.9%, **Fig. 5b** and **Supplementary Table 13**). Given the much higher number of SSVs, it was unexpected that 29 out of 30 exon-disrupting inversions were found as subcomponents of CSVs rather than standalone SSVs **(Fig. 5c)**, highlighting Swave’s ability to resolve inversion complexity. Most of exon-disrupting CSVs (30/34) altered only a single gene by modifying exon content through one or multiple CSV subcomponents. One example involved a duplicated-inversion flanked by a deletion that disrupted the 8th exon of *CYP17A1* **(Fig. 5d)**. While small variants affecting this exon have already been classified as pathogenic for congenital adrenal hyperplasia in ClinVar (**Supplementary Table 14**), this more complex rearrangement illustrates Swave’s capacity to uncover previously undetected variant forms. Annotation of the altered gene sequence using four tools revealed that the 8^th^ coding exon was incomplete compared to the original gene structure (**Fig. 5f**), leading to the loss of residues 415-508 (**Fig. 5e**), including residue 442, a critical Fe-binding site of heme group **(Extended Data Fig. 7a)**. Among the remaining four exon-disrupting CSVs, three affected paralogous genes through copy number changes. A particularly illustrative case involved *VCY*/*VCY1B* paralog genes **(Fig. 5f)**: one duplication subcomponent increased *VCY* copy number, while an inversion-duplication subcomponent simultaneously increased *VCY1B* copies. Both paralogs were copy-gained within a single CSV event, but through two distinct subcomponents.

In comparison, exon-disrupting singleton SSVs (n=888 and **Supplementary Table 12**) were structurally simpler: 613 were deletions, 274 were insertions or duplications while only one was an inversion. This singleton inversion (411bp) reversed the second exon of *HYLS1*, a gene implicated in Hydrolethalus Syndrome (**Extended Data Fig. 7b**). While most exon-disrupting singleton SSVs were small, 85 exceeded 10kb in length, with the largest one reaching 69kb. For instance, a 43kb deletion spanned introns 6-14 of *FRAS1*, a gene associated with Fraser syndrome^47,48^ (**Extended Data Fig. 7c**). Collectively, these findings demonstrated Swave’s capability to resolve intricate and previously overlooked SSV and CSV patterns within rare-disease-risk genes, providing a critical resource for downstream investigations of pathogenic mechanisms.

## Discussion

The pangenome graph currently offers the most powerful framework for integrating hundreds of high-quality genome assemblies. To address the persistent challenge of resolving SVs, particularly complex SVs, from pangenome graph embedded alleles, we developed Swave. Unlike existing pangenome-based methods that infer SV types primarily from the length differences between reference and alternative alleles, Swave leverages precise alignments and introduces projection waves to denoise repetitive sequences and support SV classification. Given the series format of waves, Swave employs a Bi-LSTM network to captures both upstream and downstream contexts, significantly improve prediction accuracy. These design innovations together underpin Swave’s effectiveness and accuracy in SV characterization.

We applied Swave to healthy cohorts, including HGSVC, HPRC and CPC, generating population-scale SV profiles that serve as valuable references for future analyses. Given the structural complexity and high polymorphism, inversions were prioritized. Remarkably, using only assemblies, Swave achieved breakpoint accuracy comparable to the consensus of multiple sequencing technologies and revealed previously unrecognized complex and polymorphic inversion patterns. The population-scale power of pangenome graphs further enable us to explore SVs in large rare-disease cohort. Applying Swave to GA4K pangenome allowed us to identify singleton SVs as potentially pathogenic ones. While experimental validation was beyond the scope of this study, many of these newly discovered variants disrupted known disease-associated genes or introduced novel alterations beyond known pathogenic variants. Considering that over 90% of GA4K samples remained undiagnosed after standard analysis of microarray and sequencing data, SV-level investigation using Swave may contribute to uncovering the mechanisms behind these previously unexplained rare-diseases.

Despite its strengths, Swave’s reliance on qualities of genome assemblies and pangenome graphs introduces some limitations. Although it demonstrated strong genotyping accuracy across population-scale samples, a subset of genotypes remained missing. These cases were attributed to incomplete assemblies, as no sequences from carrier genomes were mapped to the corresponding snarls (**Extended Data Fig.8a**). Additional analysis revealed that these problematic snarls were concentrated near centromeres and telomeres (**Extended Data Fig.8b**), suggesting that continued improvements in assembly quality, particularly in these highly repetitive regions, will be the key to close the remaining gaps.

Overall, Swave expands the utility of pangenome for comprehensive SV discovery and interpretation. Future application could leverage Swave and pangenome to enhance evolutionary analysis of SSVs and CSVs across diverse human ethnic groups or comparison between human and closely related primates, enabling the reconstruction of SV-driven evolutionary trajectories. Clinically, Swave offers a promising way for building comprehensive SV catalogs that facilitate the identification of pathogenic SVs, advancing our understanding of genetic disease mechanisms and supporting diagnosis of genetic diseases.

## Methods

### Swave methodology

#### Pangenome graph construction and allele extraction

Swave applies Minigraph for pangenome graph construction (**Extended Data Fig. 1a**), which supports identification of structural variants larger than 50 bp. Alternative tools such as PGGB^33^ and Minigraph-cactus^32^, perform more fine-grained sequence alignment and graph construction suitable for smaller variants (e.g. SNPs and Indels), which are not involved in this study. The resulting graph is encoded in GFA format (**Extended Data Fig. 1b**), comprising nodes (sequence segments with associated lengths) and edges (connections between nodes). Paths, formed by concatenating edge-linked-nodes, represent individual assembly or reference traversals through the graph.

Minigraph offers a realignment process (the ‘--call’ option) that maps each assembly back to the graph to recover its traversal paths. Graph regions where the assembly paths diverge are decomposed into snarls, which are considered as candidate SV loci (**Extended Data Fig. 1b**). For each snarl, Swave extracts traversal allele paths and reconstruct full sequences using node information in the GFA. These allele sequences are then passed to downstream modules for dotplot-based realignment and SV type classification. Allele frequencies are computed by tallying the number of carrier assemblies per variant (**Extended Data Fig. 1c**).

#### Dotplot image representation

To evaluate sequence-level difference between reference (REF) and alternative (ALT) alleles, Swave generates three types of dotplot images:

- REF2REF: alignment of reference sequence against itself, revealing the genomic baseline of REF.
- ALT2ALT: alignment of alternative sequence against itself, revealing the genomic baseline of ALT.
- REF2ALT: alignment of alternative sequence (y-axis) against the reference sequence (x-axis), revealing the structural divergence between them.

1. **Dotplot generation.** Swave begins by extracting all the kmers (default k = 30bp, stride = 1bp) from the longer sequence (REF or ALT), then traverses kmers in the shorter sequence to identify exact forward or reverse matches. For example (**Extended Data Fig. 2a above**), at the condition of *REF.length* > *ALT.length*, Swave collects all the kmers from REF sequence. For genome position *REF_i_*, the kmer with size *k* is defined as the *k* bp sub-sequence from [*REF_i_*: *REF_i+k_*). If it matches with the kmer from *ALT_j_*, Swave places a dot at the coordinate of [*REFi*, *ALTj*] in the image. The matching orientations (forward and reverse) are recorded. Then Swave move to next kmer *REF_i+stride_* and repeat the above process. After constructing the full dot matrix, Swave identifies linear dot clusters exceeding 50 bp in length (aligns with Swave’s focus on SVs), which represent structural continuity between reference and allele, serving as the skeleton of the dotplot.
2. **Dotplot optimization.** As the kmer-based approach lacks single-base resolution due to the fact that alignment gap flanking SV breakpoints may appear up to *(k - 1)* bases longer than their true lengths (**Extended Data Fig. 2a above**), Swave performs localized base-level remapping by examining whether the bases can exactly match at the kmer stop-matching boundaries (**Extended Data Fig. 2a bottom**), which are defined as the two endpoints of lines identified above. To enhance efficiency, this is applied only to lines uniquely present in the REF2ALT dotplot, excluding those shared with REF2REF or ALT2ALT dotplots, which typically reflect repetitive genomic backgrounds rather than informative SV signals.

#### Dotplot projection for waves

A major challenge in using dotplot images is the massive and redundant dots caused by genome repetitive sequences. To suppress noise from repetitive sequences and enhance SV signals, Swave introduces a novel representation form called projection waves, which distills alignment information at each genomic position from the dotplot image (**Fig.1b left**). For example, when projecting a dotplot onto its x-axis, Swave traverses each position X_i_ along the x-axis and counts the number of dots aligned across all y-axis rows (Y_1_ to Y_n_) at X_i_. Therefore, the wave summaries the matching dot number at each genome position within the projection axis. Each projection operation will generate two waves, one for forward matches and one for reverse matches. This dual-wave design preserves strand orientations, aiding in the detection of inversions.

1. **Background wave.** Swave first projects the REF2REF dotplot onto its x-axis to generate the dual-wave representing reference sequence background. Similarly, it projects the ALT2ALT dotplot to obtain alternative allele background. These background waves reflect the internal repetitive features of each allele sequence. The average value of these background waves reflects the level of repetitiveness. For example, a sequence without any repetitive components will have a wave displaying a uniform distribution with an average value close to 1 (**Extended Data Fig. 2b left**). In contrast, a sequence with dense repetitive contents produces an obviously fluctuating wave with an elevated average wave value (**Fig.1b middle** and **Extended Data Fig. 2b right**).
2. **SV-indicating wave.** Next, Swave projects the REF2ALT dotplot onto the x-axis (reference sequence) to generate the SV-indicating dual-wave (**Fig.1b right**). Dots that form the lines identified above are assigned a higher weight in the projection process, with the default weight set to the average value of the background wave. By comparing the resulting SV-indicating dual-wave to the REF2REF background dual-wave, different SV types lead to characteristic wave shifts: deletions manifest as local wave reduction, duplications as elevations, and inversions exhibit new emerging value peaks in the wave recording reversely matched dots. In addition, since insertions can be interpreted as relative deletions happened in the REF sequence, Swave also projects the REF2ALT dotplot onto the y-axis (alternative sequence). Insertions will manifest as reduced waves compared to the ALT2ALT background waves, analogous to how deletions alter the REF2REF background waves.

#### RNN for SV classification

The projection waves are stored in a series data format, and therefore, Swave leverages a RNN architecture for SV type prediction.

1. **RNN design.** Wave-derived data are encoded as sequences of four-feature tuples per reference position (**Extended Data Fig. 3a**): (i) genome position, (ii) average value of background wave, and (iii-iv) differences between SV-indicating and background waves for forward and reverse matches, respectively. To reduce redundancy, Swave merges consecutive genome positions with identical values for the latter three features, and then replaces the first feature of individual positions with the spanning lengths. Nevertheless, a single SV region may still contain multiple tuples due to the presence of genome repeats (**Extended Data Fig. 3a**). To address this, Swave employs a Bi-LSTM architecture as the core of RNN (**Extended Data Fig. 3b**), enabling the model to consider the entire context. This contextual awareness helps accurately classify multiple tuples as belonging to the same SV type.
2. **RNN training.** Swave utilizes a simulated dataset, which provide the exactly accurate SV types and breakpoints, to train the RNN mode. Real-world SV dataset are derived from callers, leading to potential inaccuracy and limited coverage of inversions and duplications. The simulated dataset comprises of five SSVs, including insertion, deletion, inversion, duplication, and inverted duplication. For each simulated SSV, its corresponding dotplot and projection waves are generated as described above, and labels for each wave tuple are directly assigned based on the coordinates of the simulated SV region.

Following the pipeline applied by SVision-pro, 1,000 events were simulated for each of the five SSV types using the VISOR randomregion.R module. Note that, for the two duplication SSVs, dispersed events and tandem events were equally simulated (500 events for each). The simulated dataset is split 70:30 into training set and validation sets. During the training procedure, the batch size is set to 128 and the learning rate is set to 0.0001. The loss function is defined as the Cross Entropy Loss. Adam Optimizer is utilized to guild the training process. Additionally, an early stopping strategy is implemented to determine the best trained models. This strategy ends the training when the validation accuracy remains unchanged (within a tolerance of 0.001) for a continuous period of 30 epochs. The training process ended at the 7^th^ epoch and in the validation set, the trained model achieved an average of 0.99 classification accuracy on the validation set.

### Benchmarking methodology

#### Assembly alignment

All genome assemblies were aligned to the human reference genome using minimap2^49^ with the following parameters ‘--MD -x asm20 -m 10000 -z 10000,50 -r 50000 --end-bonus=100 --secondary=no -O 5,56 -E 4,1 -B 5 -a --eqx -Y’. These settings were derived from previous studies^15,50^ and offered improved alignments than default configurations.

#### Pangenome graph construction

The reference genome and sample assemblies were sequentially inputted into the Minigraph with parameter ‘-cxggs’ for pangenome graph construction. To avoid naming conflicts that may generate warnings in Minigraph during graph construction, all contigs were renamed to include sample and haplotype identifiers. Following construction, the input sequences were mapped back to the graph and the corresponding paths for each graph snarls were determined by Minigraph --call function. Full assembly-specific paths through the graph were then reconstructed using a custom script that concatenates individual snarl paths sorted by reference genome positions.

#### Individual-level benchmark

For SSV detection, the HG002 assembly and Tire 1 high-confidence SVs set, and human genome hg19 were used as ground truth. PAV and SVIM-asm were run with default parameters. For vg-vcfwave, the pipeline was initialized with ‘vg deconstruct’ to produce a VCF file recording the snarl and path information. Vcfbub and vcfwave were then executed following the released script by HPRC. For CSV detection, the same ground-truth dataset used in the SVision-pro study was adopted. Swave called CSVs from the pangenome graph built with human reference hg38 and template genome with all CSVs inserted. The assembly-based SVision-pro callset was obtained by applying SVision-pro on the alignment results between temple genome and hg38 reference genome. The read-based SVision-pro callset was obtained from its publication. For both SSVs and CSVs, performance was assessed using Truvari, calculating F1-score against the corresponding ground truth.

#### Trio-level benchmark

Pangenome graphs were constructed for each trio along with human reference genome T2Tv2.0. All merge tools (Jasmine, SURVIVOR and Truvari), and the refinement tool PanPop, were executed following their official instruction to obtain the integrated trio callset including genotypes for each trio member. Swave and vg-vcfwave could directly output the callset comprising genotypes for each individual. To calculate the mendelian consistency, we first collected the genotypes of parents and generated the list of all possible child genotypes. If the actual child genotype matched with anyone in the list, we determined it as a consistent one.

#### Population-level benchmark

To assess genotyping accuracy at the population level, pangenome graphs were built using reference genome T2Tv2.0 and 130 haplotypes from HGSVC. Callers were executed as the same as in the trio-level benchmark. Genotype completeness was evaluated at the haplotype-level by measuring the proportion of missing haplotype (denoted as ‘.’) across all SV records.

#### Computational inversion validation

To computationally validation inversions, we reconstructed the inversion-feature sequence by reversing the reference corresponding to each detected inversion region. The sequence was then mapped to the carrier haplotype using mappy, a python interface to minimap2. Carrier haplotype sequence was reconstructed by concatenating sequences of nodes from the pangenome graph corresponding to the haplotype path. If the detected inversion is true-positive, the inversion-feature sequence would forwardly and continuously map to the carrier haplotype sequence with no clips. Therefore, mapping integrity was defined as the ratio of aligned length to total inversion-feature length. We also leveraged the validation metrics from two published tools, TT-mars and Vapor. Both the two tools measure whether the inversion-feature sequence exhibits a better match to the carrier haplotype than the wild-type reference sequence, with TT-mars using an aligner (mappy) while Vapor using dotplot to get the mapping results. Their core validation modules were integrated into the overall mapping integrality framework mentioned above. For CSVs with multiple inversion subcomponents, each inversion substructure was validated independently.

#### Gene and repeat annotations

Gene annotation was performed using ANNOVAR^51^ following the official instructions. For CSVs, each subcomponent was annotated separately. For duplications, both source and inserted regions were annotated. SVs intersecting ‘exons’ or ‘splicing’ annotations were flagged for downstream analysis. Repeat annotation was performed using Tandem Repeat Finder (TRF)^52^ with recommended parameters on both the reference and alternative sequences. A repeat ratio was calculated for each allele as the fraction of repeat-annotated bases over total length. SVs were classified as in “highly-repetitive regions”, if the maximum repeat ratio of either reference and alternative sequence annotation exceeded 80%.

## Data availability

All the published reference genomes, sample assemblies, and SV callsets are listed in **Supplementary Table 15**. The callsets on healthy and disease cohorts produced by Swave are shared in Google Drive (https://drive.google.com/drive/folders/1ibqKfIyQe_ErpFi-v6oEG9AhlsD3EUC7?usp=drive_link)

## Code availability

Swave (v1.0) is available at GitHub (https://github.com/songbowang125/Swave.git). The scripts for reproducing the results in this paper are available at GitHub (https://github.com/songbowang125/Swave-Utils.git)

## Acknowledgements

K.Y. is supported by the National Key R&D Program of China (grant no. 2022YFC3400300) and National Science Foundation of China (grant nos. 32125009 and 32430017). S.W. is supported by National Science Foundation of China (grant nos. 323B2015)

## Author contributions

K.Y. designed and supervised the research. S.W. developed the algorithm and performed the performance evaluation and downstream analysis. T.X. and P.Z. analyzed the impact of SVs.

## Extended Figures

**Extended Data Fig. 1.**
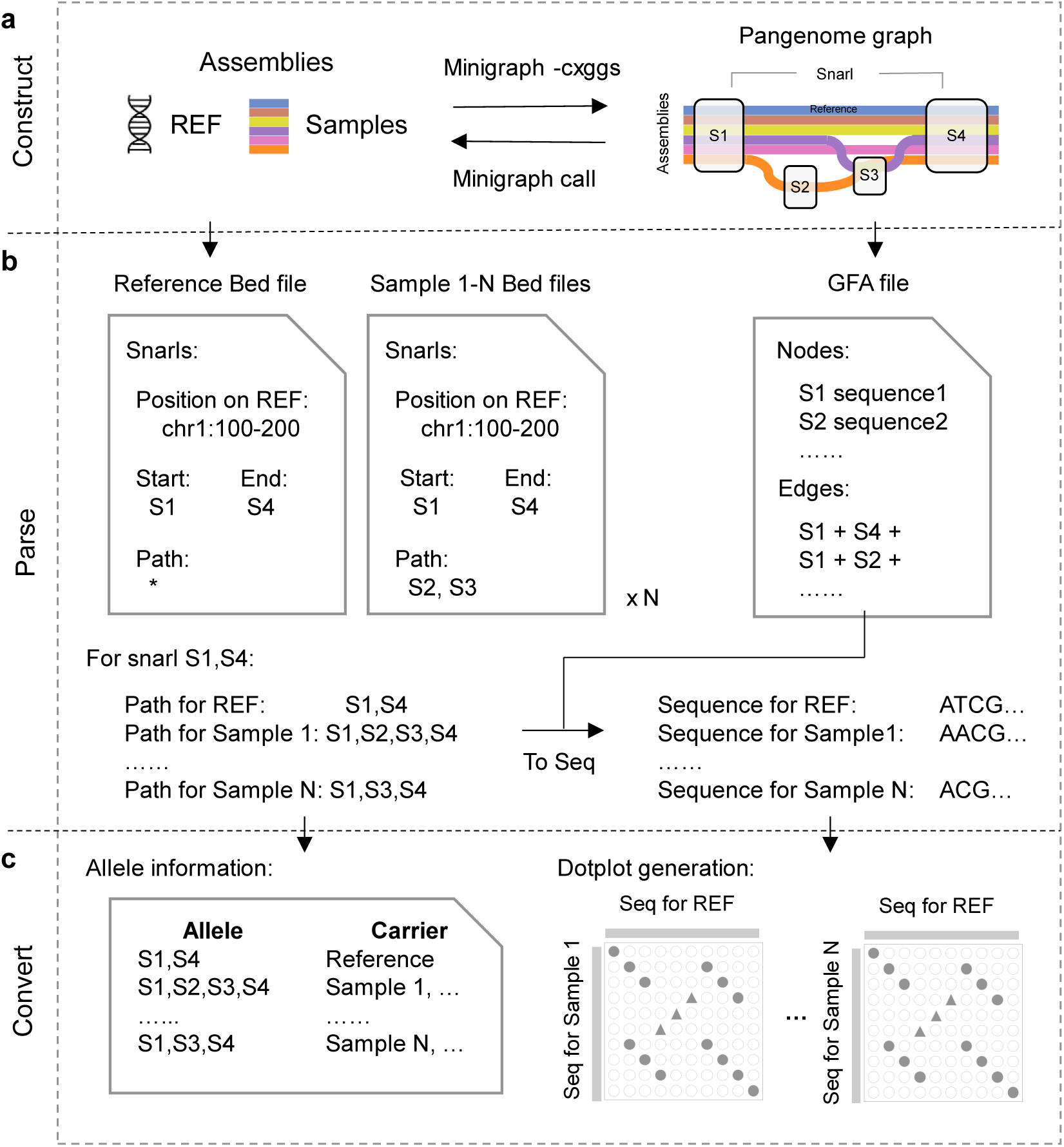
Overview of pangenome construction and allele extraction in Swave. **a,** Construction of pangenome graph using Minigraph with both reference and sample assemblies. The resulting graph is saved in GFA format, which encodes node sequences and directed edges between nodes. **b,** Assembly paths are recovered using --call function in Minigraph. Regions where paths diverge (Snarls) are identified as candidate structural variant loci. Allele sequences for each snarl are reconstructed by extracting the corresponding node sequences from the GFA. **c,** Based on the Minigraph --all outputs, Swave determines carrier assemblies for each allele and proceeds to generate dotplots for each reference-alternative pair in the next processing module.

**Extended Data Fig. 2.**
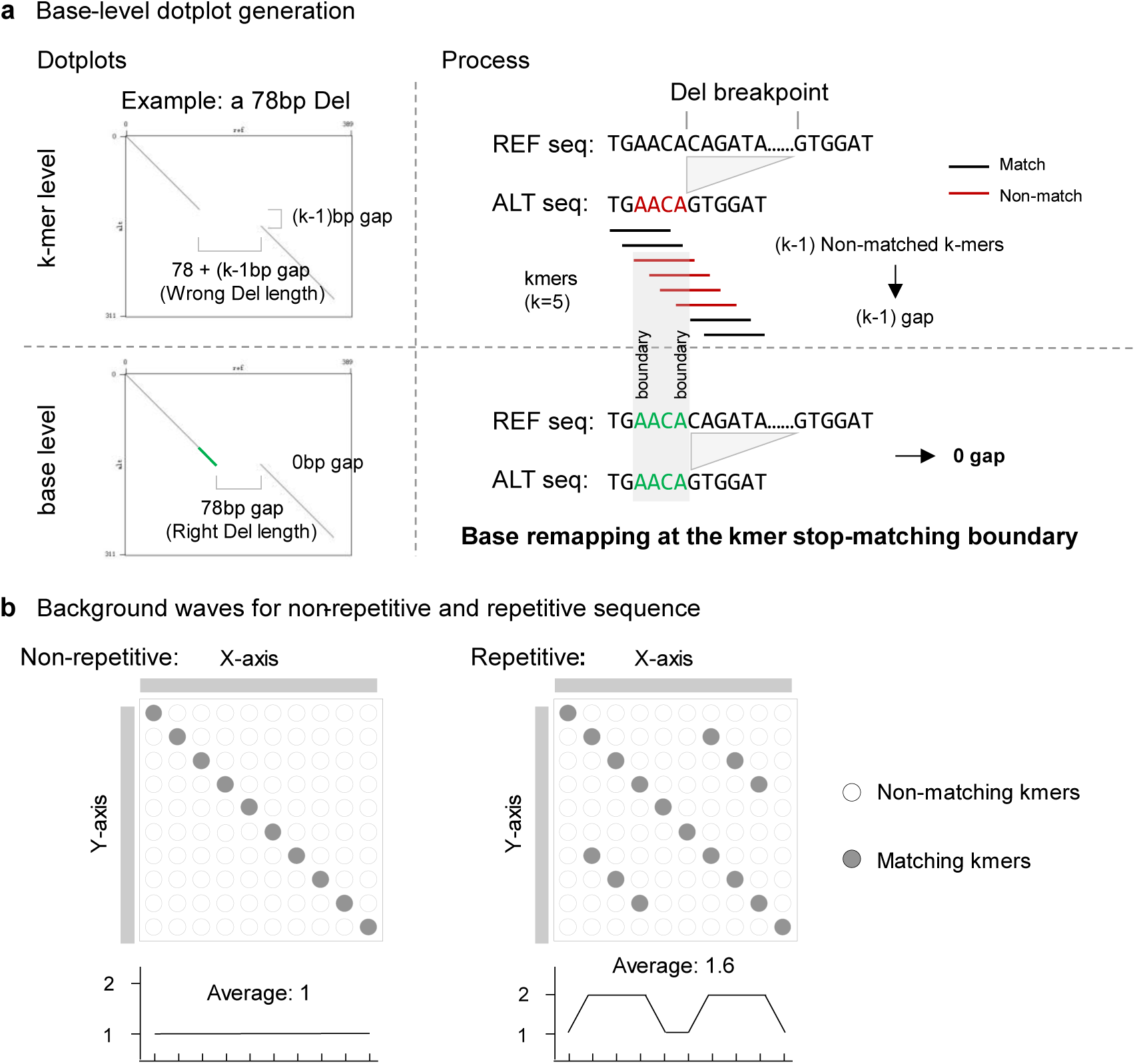
Dotplot generation and projection. **a,** Base-level refinement of kmer-based dotplots. Initial alignment introduces (k-1) base gaps near SV breakpoints. Swave performs base-level remapping at kmer stop-matching boundaries to improve breakpoint resolution for downstream SV classification. **b,** Influence of genomic repeats on wave patterns. Dense, repetitive regions generate abundant spurious matches in dotplots, resulting fluctuating wave signals upon projection.

**Extended Data Fig. 3.**
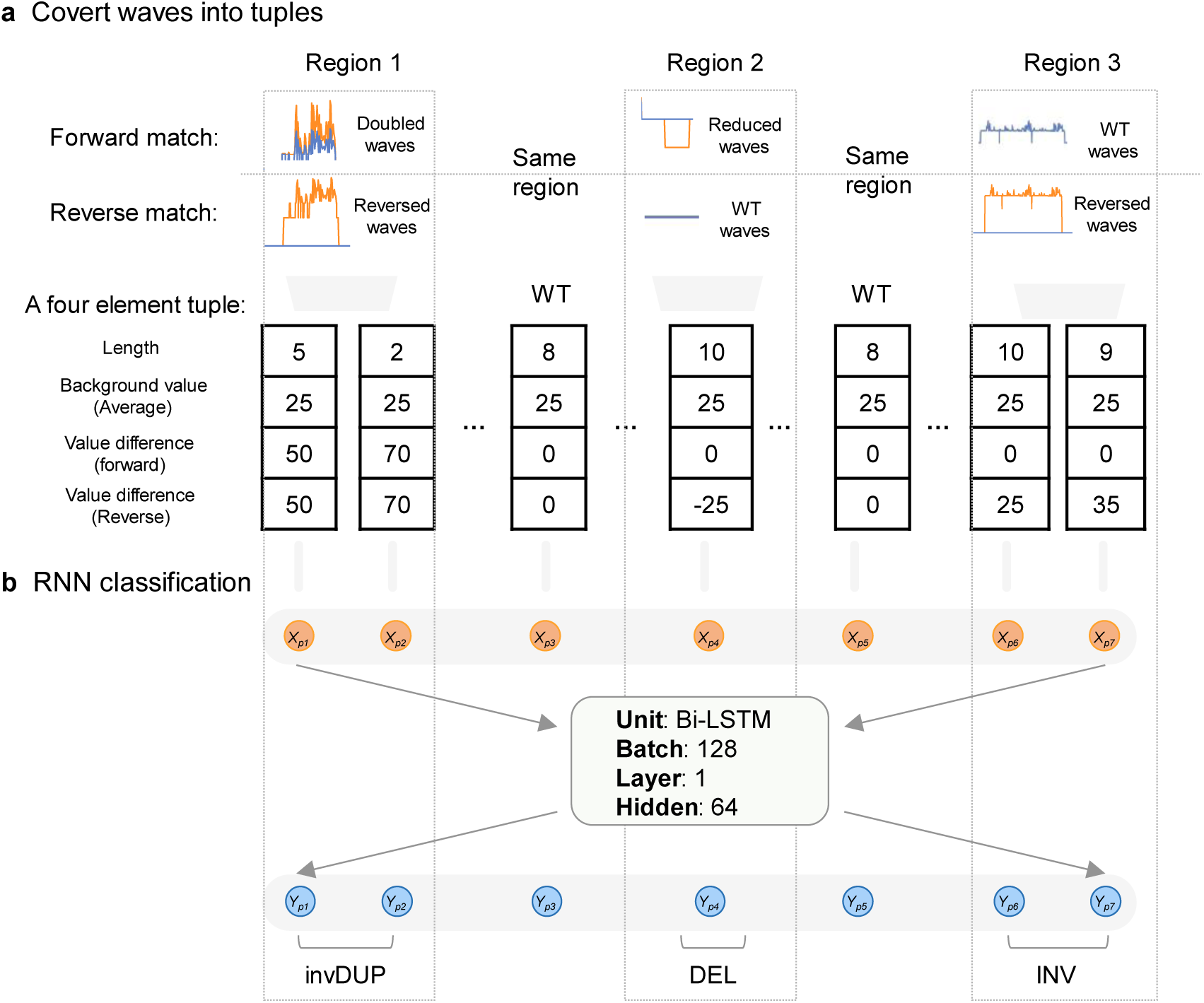
Recurrent Neural Network for SV classification in Swave. **a,** Projected wave signals are encoded as four-element tuples per genomic segment, comprising span length, background average wave value, and the differences between SV-implying and background waves for both forward and reverse matches. These tuples serve as the input for the RNN classification model. **b,** A one-layer Bi-LSTM with 64 hidden units forms the core of the RNN, enabling context-aware classification of SV components across the sequence.

**Extended Data Fig. 4.**
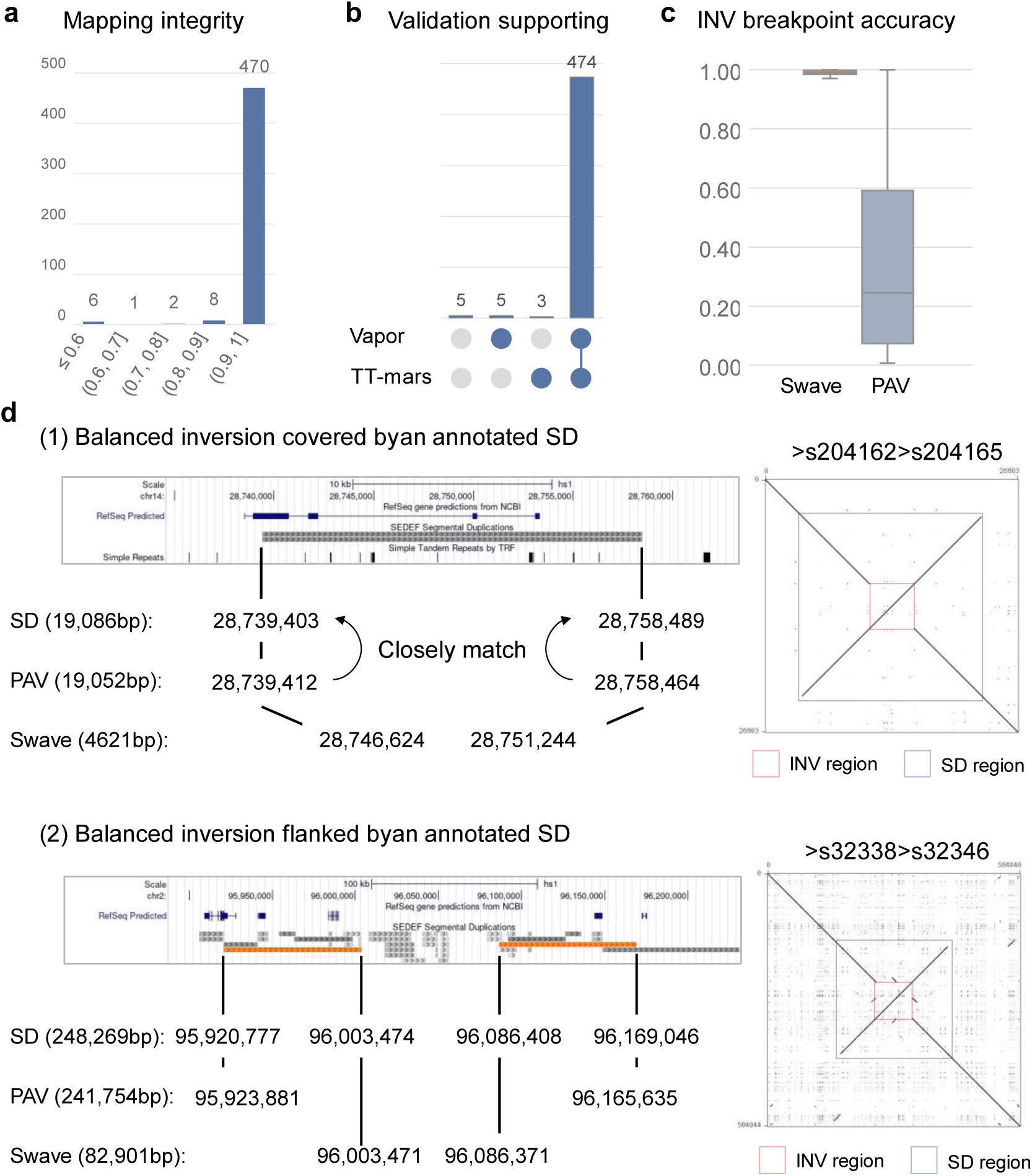
Validation and illustration of inversions. **a and b,** Validation results for all detected balanced and complex inversions. Three orthogonal metrics were applied: mapping integrity, TT-mars, and Vapor. **c,** Comparison of inversion breakpoint accuracy between Swave and PAV. Bounds represent the interquartile range (IQR) from Q1 to Q3. Whiskers extend to values within Q1 − 1.5*IQR and Q3 + 1.5*IQR. Minimum and maximum values are shown. **d,** Illustration of breakpoint distortion caused by inverted segmental duplications (SDs). While PAV’s breakpoints are frequently shifted due to alignment ambiguity, Swave maintains accurate breakpoint placement within repetitive regions.

**Extended Data Fig. 5.**
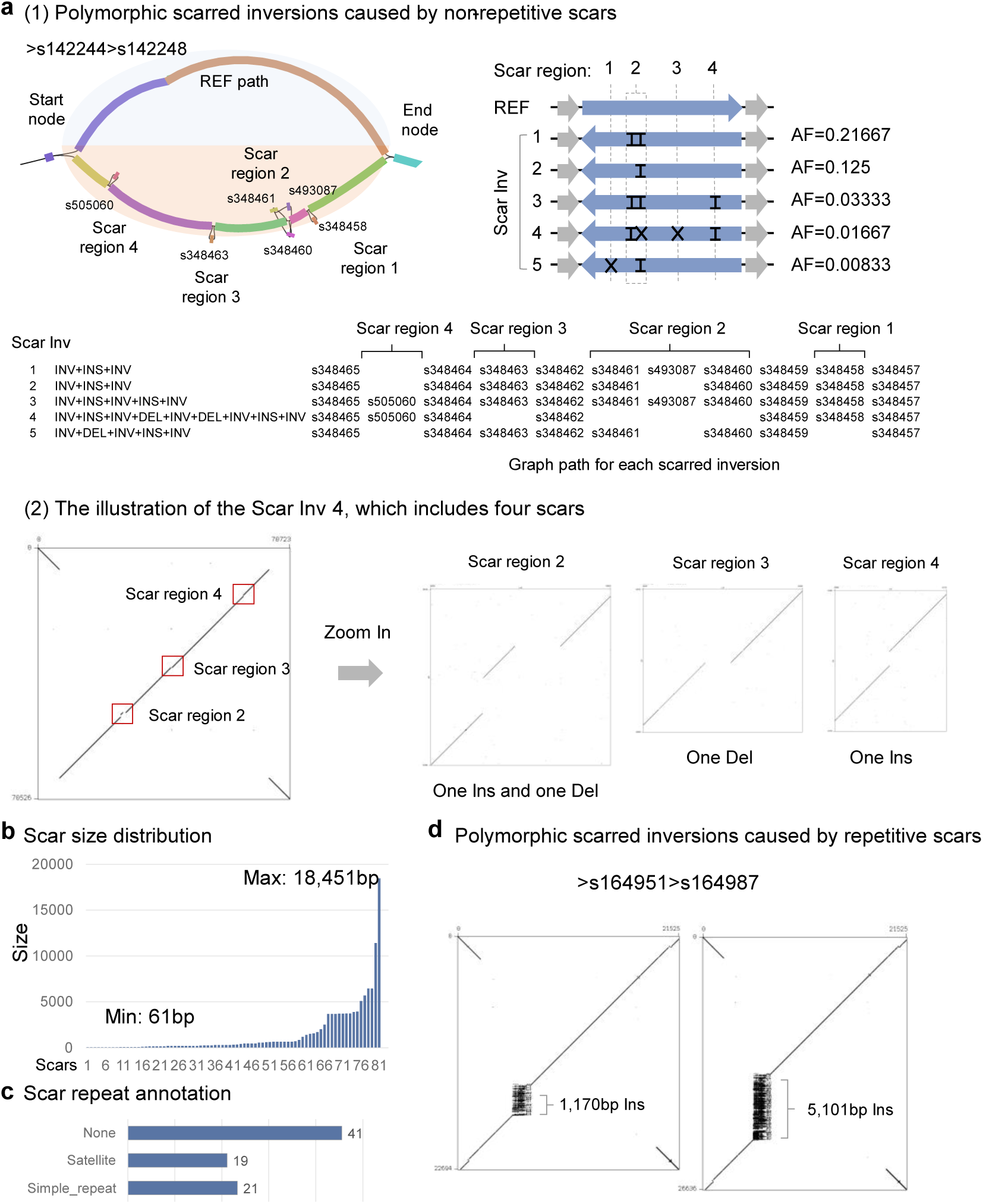
Characterization of polymorphic scarred inversions. **a,** Example of a polymorphic scarred inversion snarl containing five distinct alleles (1), generated by combinatorial arrangements of five unique internal scars across four genomic regions. The most complex variant includes four separate scars (2). **b,** Length distribution of all detected scars (n=81), ranging from 61bp to 18,451bp. **c,** Repeat annotation of all scars (n=81). **d,** Example of polymorphic scarred inversions driven by repetitive elements, where two repeat expansions give rise to insertion scars of difference lengths.

**Extended Data Fig. 6.**
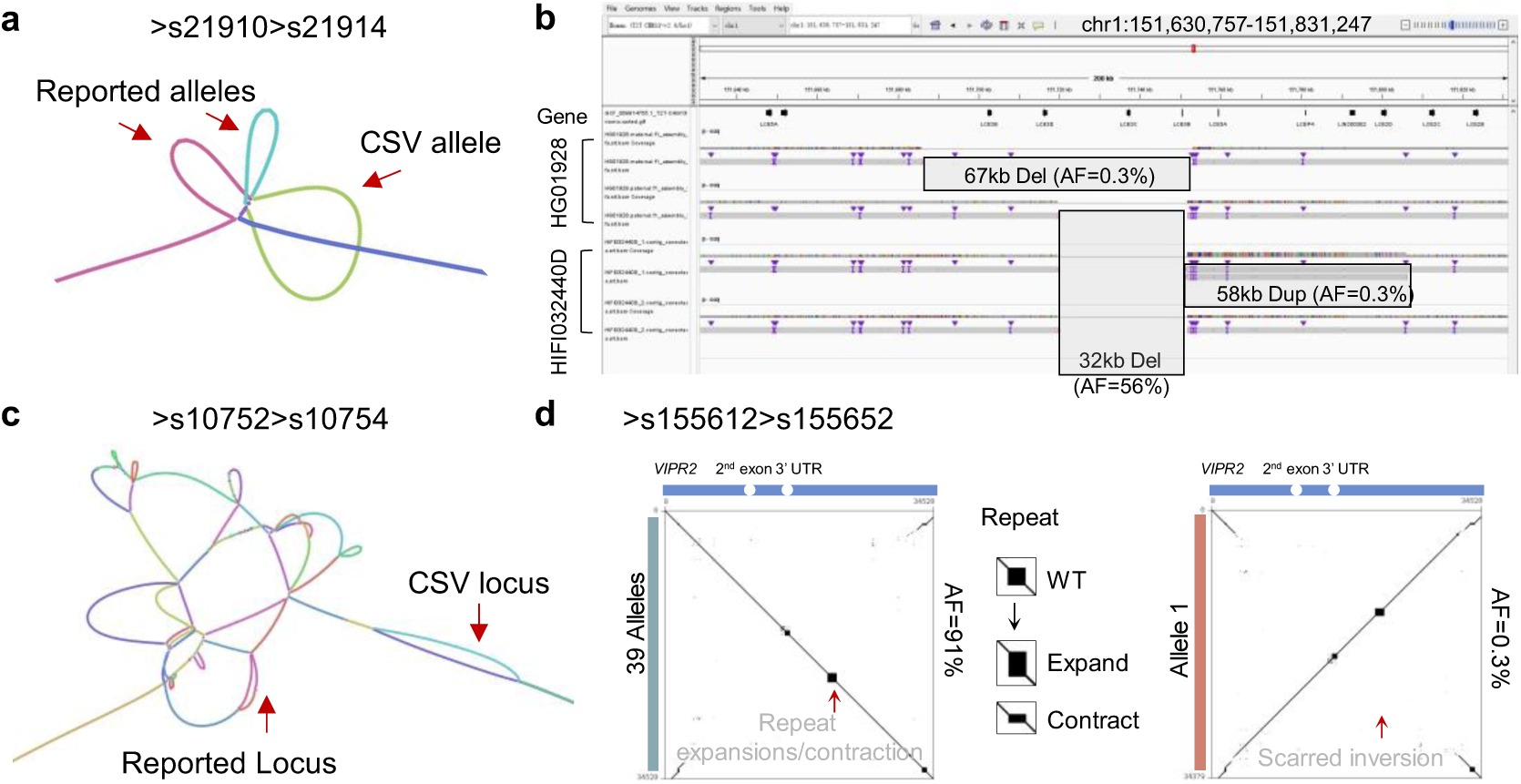
Rare and complex structural variants revealed by Swave. **a,** Pangenome graph structure of snarl ‘>s21910>s21914’. A rare CSV allele introduced a novel traversal path not observed among the reported alleles. **b,** IGV snapshot of snarl ‘>s21910>s21914’, illustrating co-occurrence of two distinct SSVs and one CSV. The rare SSV (67kb deletion) extended the common 32bp deletion, where the rare CSV (a duplication flanked by a deletion) added a 58kb duplication at the right breakpoint of the frequent 32kb simple deletion. **c,** Pangenome graph of snarl ‘>s10752>s10754’, showing a novel CSV locus, structurally distinct from previously reported variants. **d,** Illustration of a rare scarred inversion that partially disrupts the coding structure of *VIPR2*, a gene associated with neuropsychiatric disorder.

**Extended Data Fig. 7.**
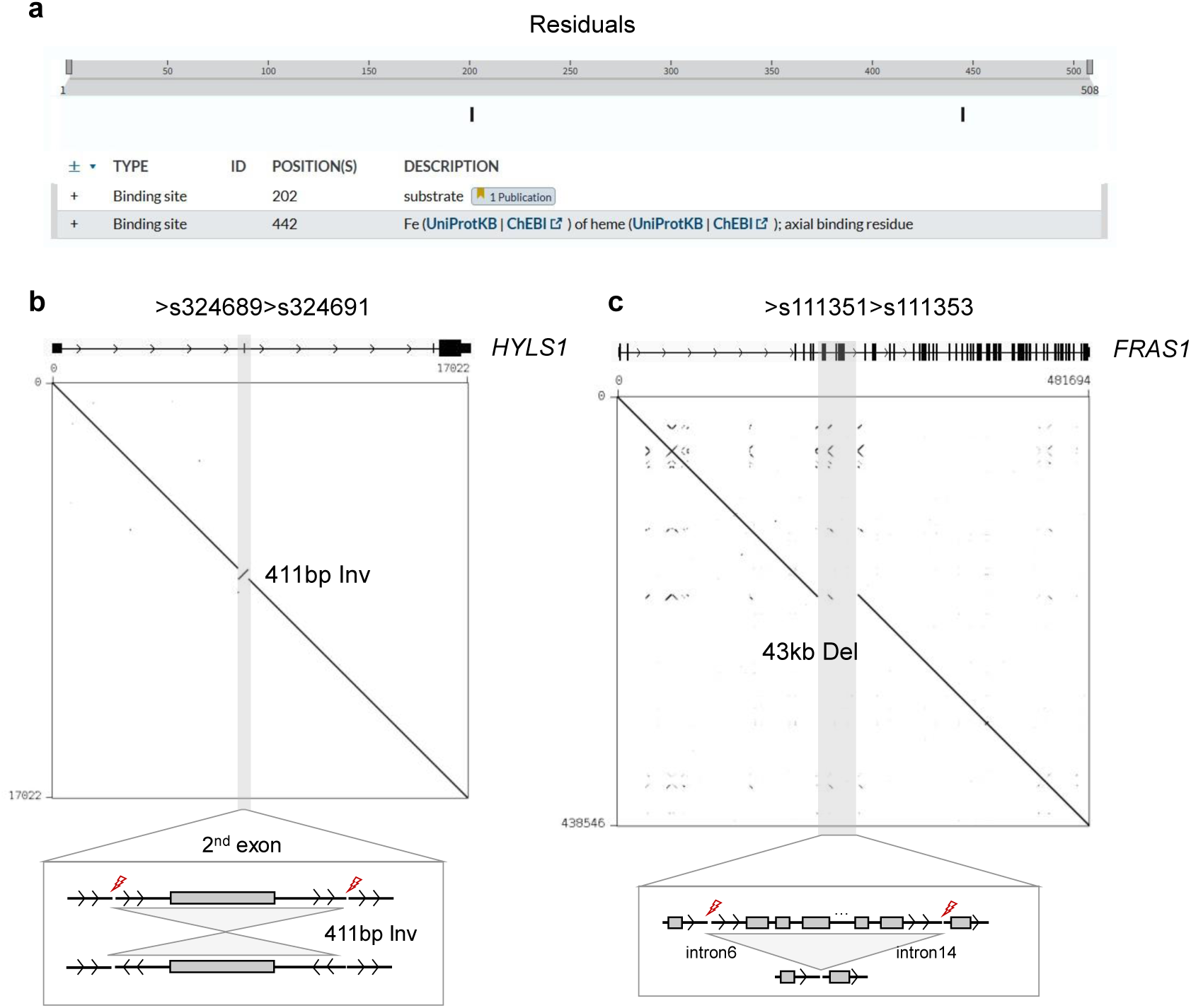
**a,** Structural annotation of the CYP17A1 protein highlights two functional binding sites, as sourced from UniProt**. b,** Schematic of a simple structural variant, a 411bp inversion, disrupting the 2^nd^ exon of gene *HYLS1*, a gene implicated in Hydrolethalus Syndrome. **c,** Representation of a 43kb deletion spanning introns 6 to 14 of gene *FRAS1*, a gene associated with Fraser syndrome.

**Extended Data Fig. 8.**
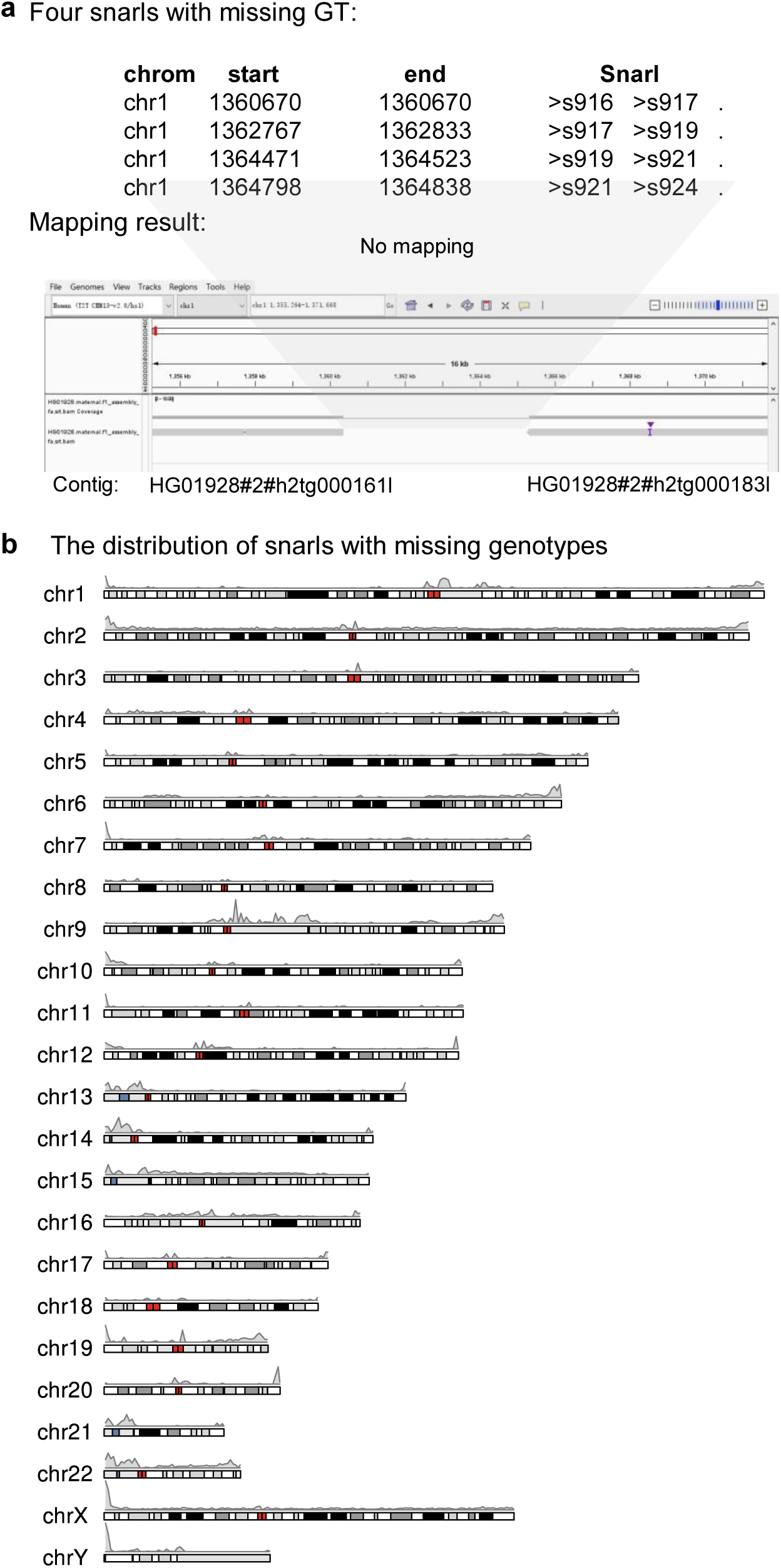
Genotyping incompleteness associated with unresolved pangenome graph regions. **a,** Mapping results of a carrier assembly exhibiting missing genotypes across four snarls. **b,** Genome-wide distribution of snarls with missing genotypes across HGSVC samples. The Y-axis indicates the number of assemblies lacking mappable sequence at each snarl.

